# Motor Unit Number Estimation Using Magnetomyography

**DOI:** 10.64898/2026.05.25.727665

**Authors:** Burak Senay, Nima Noury, Markus Siegel, Oliver Röhrle, Thomas Klotz, Justus Marquetand

## Abstract

**Objective:** Investigation of the feasibility and characteristics of contactless motor unit number estimation (MUNE) using optically pumped magnetometer–based magnetomyography (OPM-MMG) as compared to surface electromyography (EMG).

**Methods:** Simultaneous electrically evoked OPM-MMG and EMG signals of the abductor digiti minimi muscle (ADM) were measured in three healthy participants. To characterize MUNE across both modalities and account for within-subject physiological variability, 20 repetitions of electrical stimulation of the ulnar nerve at randomized intensities ranging from 5 to 30 mA in 0.1 mA increments were performed, resulting in a total of 5,020 evoked responses per subject. We quantitatively compared of MUNE and evoked EMG/MMG signal characteristics, including signal-to-noise ratio (SNR) and motor unit response amplitudes. SNR was equalized between measurement modalities to estimate the effect of SNR on MUNE.

**Results:** MMG-derived MUNE (mMUNE) could be assessed contactlessly. mMUNE estimates were on average 40% lower than EMG-derived MUNE (eMUNE), ranging from 30–65 for mMUNE versus 69–101 for eMUNE. Equalizing the EMG SNR (29–31 dB) to match the MMG SNR (18–27 dB) yielded comparable eMUNE and mMUNE estimates. Peak-to-peak amplitudes of the supramaximal compound motor unit fields ranged from 34–73 pT and single motor unit fields ranged from 0.7–1.4 pT.

**Significance:** These findings demonstrate that OPM-MMG enables contactless motor unit number estimation.

## Introduction

Motor units, consisting of a single alpha motor neuron and all the muscle fibers it innervates, represent the fundamental functional units of the neuromuscular system (Sherrington, 1919). The number of motor units within a muscle is closely linked to force production, motor control, and adaptive or pathological changes in neuromuscular organization. Reductions in motor unit number have been reported with advancing age and in several neuromuscular disorders involving denervation and reinnervation processes, including motor neuron and peripheral neuromuscular diseases (Gooch et al., 2009; Hansen and Ballantyne, 1978, p. 1; McComas et al., 1971; Power et al., 2013). These observations motivate the use of the motor unit number as an indirect marker of neuromuscular integrity. Because direct counting of motor units is currently not feasible in living humans, indirect methods have been developed to estimate motor unit numbers and are typically referred to as MUNE.

Early MUNE methods were introduced several decades ago to estimate motor unit numbers from electrophysiological recordings obtained during incremental peripheral nerve stimulation (McComas et al., 1971). These methods were based on the observation that the electrical response of a whole muscle—the compound muscle action potential (CMAP)—can be interpreted as the summation of contributions from individual motor units. By estimating the average electrical contribution of a single motor unit, typically referred to as the single motor unit potential or surface motor unit potential (SMUP), the total number of motor units can be approximated as the ratio of supramaximal CMAP amplitude to mean SMUP amplitude. Some MUNE approaches rely on voluntary contractions combined with intramuscular recordings to isolate the discharge of individual motor units, which are then used as triggers to extract surface motor unit potentials through spike-triggered averaging. These methods include classical spike-triggered averaging (STA), decomposition-enhanced spike-triggered averaging (DE-STA), and related quantitative EMG approaches that sample single motor unit potentials during voluntary activation (Boe et al., 2004; Brown et al., 1988; Doherty and Stashuk, 2003; McNeil et al., 2005). Conversely, other MUNE techniques are based on incrementally electrically evoked muscle responses recorded at the skin surface during peripheral nerve stimulation. In these approaches, progressively increasing stimulation intensities recruit motor units in a stepwise manner, allowing estimation of single motor unit potentials from discrete increments in the compound muscle action potential or from modelling the stimulus–response relationship of the entire motor unit pool (Bostock, 2016; Chen et al., 2023; Doherty and Brown, 1993; McComas et al., 1971).

Although MUNE is well-established and has been performed predominantly using surface EMG, its implementation is partly constrained by several practical and methodological limitations of the recording modality. Surface EMG provides direct access to muscle electrical activity but requires physical contact with the skin, typically involving skin preparation, electrode placement, and, in some cases, invasive needle recordings. These requirements can be time-consuming, potentially uncomfortable, or impractical in certain populations, such as children. Moreover, EMG recordings are sensitive to electrode–skin impedance and local tissue properties, such as subcutaneous fat, which influence amplitude-based measurements central to MUNE calculations (Farina et al., 2010; Fuglsang-Frederiksen, 2006).

Recently, novel quantum sensors have renewed interest in contactless magnetic muscle recordings, offering an alternative approach for studying neuromuscular physiology that eliminates the need for electrode–skin contact. Magnetomyography (MMG) measures magnetic fields generated by intracellular and extracellular currents associated with muscle fiber action potentials, which, within the limits of the quasi-static approximation, can be described by the Biot–Savart law. Although MMG was proposed several decades ago (Cohen and Givler, 1972), its broader application has historically been limited by the technical constraints of available sensors. In particular, superconducting quantum interference devices (SQUIDs), although highly sensitive, require cryogenic cooling and rigid sensor geometries, which restrict their use to well-defined body regions (e.g., helmets for brain recordings) and static conditions. This limitation of rigid sensor geometries was overcome in the past decade, when non-cryogenic, small and modular sensors – optically pumped magnetometer (OPMs), with sensitivities as low as ∼15 fT/√Hz – became commercially available. OPMs can be positioned flexibly with respect to the muscle of interest, enabling individualized sensor configurations without physical contact (Boto et al., 2017; Kominis et al., 2003). It was demonstrated that OPM-based MMG (OPM-MMG) can measure neuromuscular signals with temporal and spectral characteristics highly similar to those observed in EMG (r > 0.9), while potentially offering advantages in comfort, price, experimental flexibility, and measurement setup compared to EMG (Broser et al., 2021; Marquetand et al., 2021).

These developments naturally raise the question of whether established MUNE in EMG can be translated to contactless OPM-based magnetomyography, i.e., if MMG-derived MUNE (mMUNE) is feasible and how it compares to EMG-derived MUNE (eMUNE). Beyond feasibility, addressing these questions allows us to quantify sensor requirements *in vivo*, thereby informing future sensor development.

To address these questions, we simultaneously recorded electrically evoked surface EMG and OPM-based MMG from the abductor digiti minimi (ADM) muscle during incremental stimulation of the ulnar nerve. Using identical analysis procedures across modalities, we applied an incremental MUNE framework to both EMG and MMG recordings to estimate compound muscle responses and average single motor unit contributions. This approach enabled a direct comparison between conventional EMG-derived MUNE and the proposed MMG-derived MUNE, allowing us to evaluate whether established electrophysiological MUNE principles can be transferred to contactless magnetic measurements.

## 2. Methods

### 2.1. Participants

A total of three healthy male adults (27–28 years old) participated in the experiment. The sample size was selected primarily to demonstrate methodological feasibility and, secondarily, to characterize mMUNE relative to eMUNE. A larger sample size was not pursued, as MUNE is an individual diagnostic measure and a highly reliable metric across time points (Kaya et al., 2014); therefore, including additional subjects was not expected to yield substantially different results. To facilitate comparison between mMUNE and eMUNE, we determined that evaluating the amount of information per subject provided a more informative analysis than merely increasing the overall number of subjects in the group (see also the protocol section below). All participants provided informed consent to participate in the study and to have their data published. The local ethics committee reviewed and approved the study protocol in accordance with the World Medical Association’s Declaration of Helsinki (97/2021BO2).

### 2.2. Setup, Sensors, and Electrodes

All experiments were conducted in a magnetically shielded room (AK3b, VAC Vacuumschmelze) at the MEG Center of the University of Tübingen, Germany. Given that OPM-MMG can be measured contactlessly and that the MMG signal decays with the inverse square of the distance (Yang et al., 2025b), we explored, as an additional research question, whether the MUNE could still be accurately calculated from distances greater than 2 cm. To investigate this, we designed and created a 3D-printed planar grid consisting of 4 by 4 OPM sensors, which was positioned adjacent to the subject’s right abductor digiti minimi muscle. In total, sixteen OPMs (FieldLine Medical, V3-uniaxial, Boulder, Colorado, USA) were used to measure OPM-MMG, with each vapor cell separated by 20 mm from the others (see Figure 1A). This 4x4-OPM grid was placed as close as possible. i.e., at a maximum distance of 5 mm from the skin surface.

**Figure 1:**
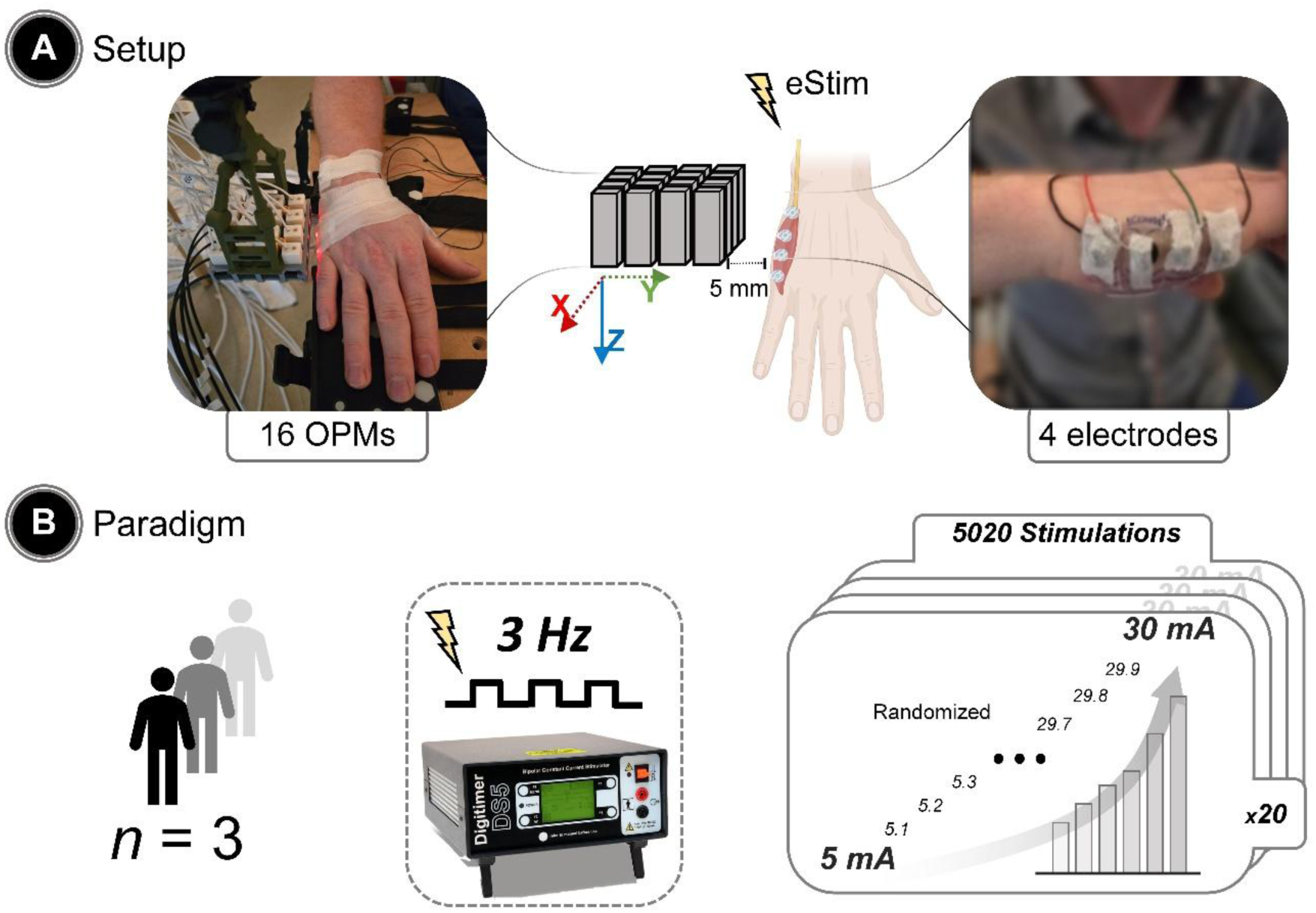
Experimental setup and paradigm. **(A)** Experimental setup with a 4×4 array of 16 OPM sensors positioned over the abductor digiti minimi (ADM) muscle and surface EMG electrodes placed on the muscle belly. **(B)** Experimental paradigm showing stepwise electrical stimulation of the ulnar nerve from 5 to 30 mA in 0.1 mA increments with 20 repetitions per intensity, delivered in randomized order with a jitter at 3 Hz. EMG and OPM-MMG signals were recorded simultaneously.

Regarding EMG, the skin over the ADM muscle was prepared by cleaning with alcohol pads and mildly abrading with a surgical scrub to ensure optimal contact. Four monopolar EMG electrodes (Conmed, Cleartrace2 MR-ECG-electrodes) were placed along the muscle belly of ADM, while ensuring visually that each electrode was placed below one OPM (see Figure 1A). These disposable wet non-magnetic carbon electrodes have a conductive circular area with a diameter of 25 mm. Because four electrodes placed adjacently would extend beyond the anatomical boundaries of the ADM muscle, the adhesive foam surrounding the conductive area was trimmed prior to placement, resulting in an effective electrode diameter of approximately 20 mm while preserving the central conductive carbon contact (10 mm diameter). The center-to-center distance between adjacent electrodes was approximately 15 mm, allowing four electrodes to cover the full extent of the ADM muscle.

Electrode placement was visually aligned with the ADM muscle belly and oriented approximately along its longitudinal axis. Each EMG electrode was carefully positioned directly underneath one of the OPM sensors to facilitate spatial correspondence between electrical and magnetic recordings. A reference electrode was placed over the styloid process of the ulna, and a ground electrode was placed at the metacarpal bone of the thumb. EMG signals were recorded in a monopolar configuration relative to this reference electrode. EMG electrodes had impedances ranging from 2 to 25 kΩ across all subjects. To ensure measurement stability during OPM-MMG recordings, the participant’s right hand was firmly fixed using a custom-built wooden support.

EMG was recorded using the EEG amplifier of the available CTF-MEG system (CTF Omega 275, Coquitlam, BC, Canada) with a sampling rate of 5860 Hz, and OPM-MMG was recorded using the manufacturer-provided data acquisition system (HEDscan, FieldLine Medical, Boulder, Colorado, USA) with a sampling rate of 2500 Hz. Electrical stimulation of the ulnar nerve was delivered using a Digitimer DS5 (Isolated Bipolar Constant Current Stimulator, UK). All data (EMG, OPM-MMG, stimulation) were synchronized via a custom MATLAB script that controlled a cDAQ-9178 data acquisition device (National Instruments, USA), which generated hardware triggers for both controlling the electrical stimulation and recording systems.

### 2.3. Protocol

To ensure recordings with both, supramaximal and minimal stimulation intensities, we applied electrical stimulation ranging from 5 to 30 mA in 0.1 mA increments, with a pulse width of 0.2 ms at a jittered 3 Hz frequency (the jitter was uniformly distributed and symmetric around 0, with a ±50 ms range). We repeated each intensity 20 times in a randomized order, resulting in 5020 evoked responses per subject (see Figure 1B).

### 2.4. Data Analysis and Processing

All data analyses were performed in MATLAB (MathWorks, R2023a) using custom scripts and the open-source FieldTrip toolbox (Oostenveld et al., 2011). The complete processing workflow is illustrated in Figure 2. Briefly, time-synchronized (see Section 2.2) OPM-MMG and surface EMG recordings were processed using identical analysis steps. Before filtering, surface EMG signals, originally recorded at 5860 Hz, were resampled to 2500 Hz using the FieldTrip toolbox to match the sampling rate of the OPM-MMG recordings. As shown in Figure 2 (step 1), raw OPM-MMG and EMG signals were filtered using least-square finite-impulse-response (FIR) filters applied in a zero-phase forward–reverse procedure. For all modalities, signals were high-pass filtered at 10 Hz and low-pass filtered at 185 Hz. Low-pass and high-pass filter frequencies were chosen to match the effective hardware bandwidth of the OPM sensors; applying the same filtering to EMG ensured that subsequent analyses were not biased by modality-specific frequency content. Narrowband Hamming window-based FIR notch filters (half-bandwidth: 1 Hz) were applied at 50, 100, 150, and 200 Hz to attenuate line-noise contamination and its harmonics. FIR filters were implemented using a least-squares design (firls function) for high-pass and low-pass filtering, where fs denotes the sampling rate (2500 Hz) and fc the cutoff frequency (10 Hz for high-pass and 185 Hz for low-pass). Transition bandwidths were defined as 20% of the cutoff frequency. Filter orders were 2500 for the high-pass filter and approximately 150 for the low-pass filter. Notch filters were implemented using fir1 function with a ±1 Hz bandwidth around 50 Hz and its harmonics (100, 150, and 200 Hz), with order scaling factors between 100 and 180.

**Figure 2:**
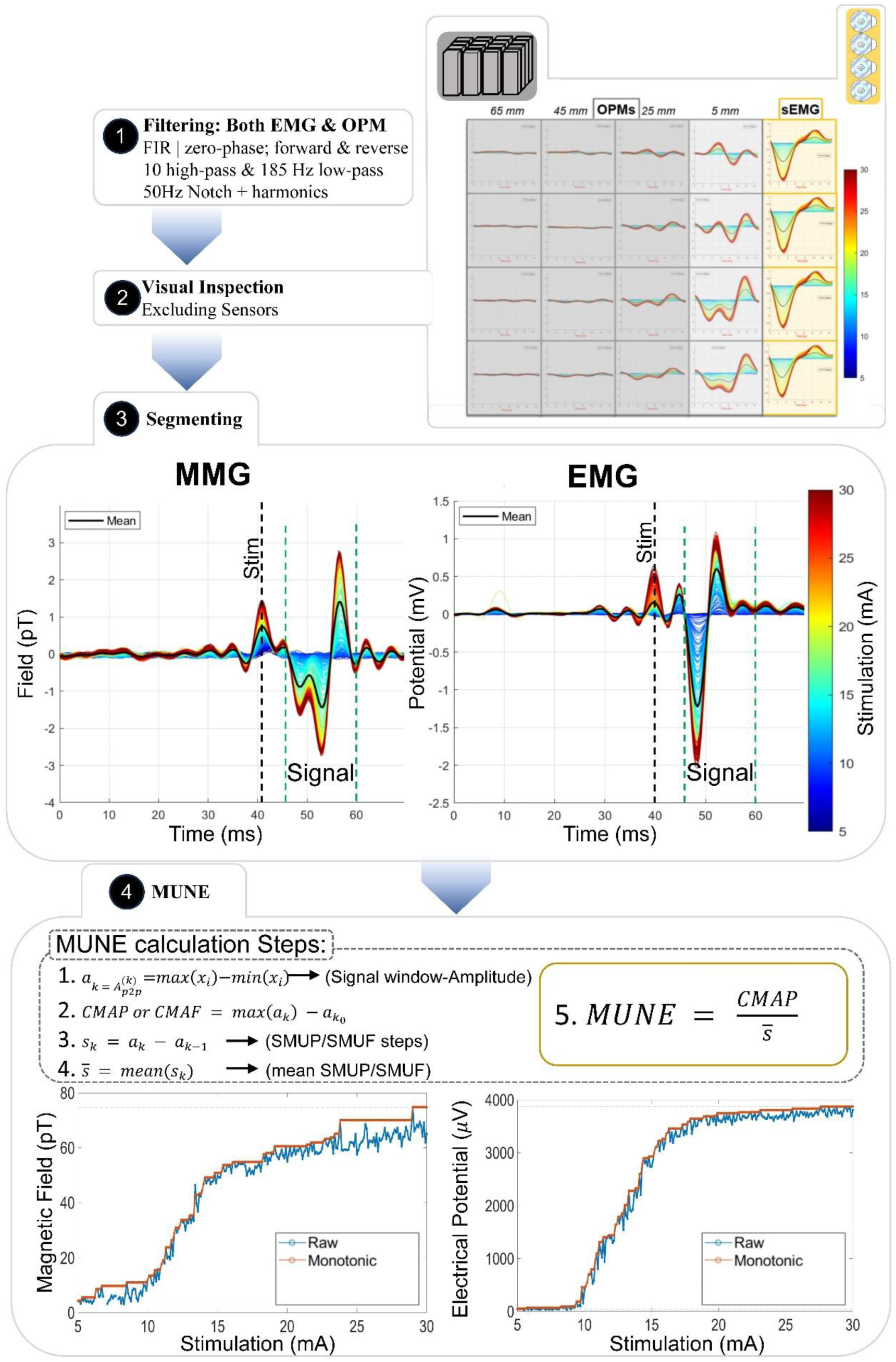
Data analysis pipeline in four main steps: **Step 1 (Filtering):** Raw sensor signals were band-pass filtered (10–185 Hz) and notch-filtered at 50 Hz and harmonics to suppress line noise and low-frequency drifts (see Methods). **Step 2 (Signal inspection and sensor selection):** After filtering, signals were visually inspected to exclude sensors with excessive noise or a lack of a clear stimulus-locked response. Example segmented signal traces of OPM-MMG are shown for increasing sensor–skin distances (top right), illustrating signal loss with distance. **Step 3 (Segmentation):** Stimulus-locked epochs were extracted from the continuous recordings. For each stimulation intensity, signal and noise windows were defined relative to the stimulation onset (dashed and dotted lines), yielding segmented MMG and sEMG responses across repetitions**. Step 4 (MUNE computation):** Response amplitudes were quantified within the signal window using the peak-to-peak metric. Amplitude–intensity curves were constructed per repetition, converted to non-decreasing recruitment curves, and used to compute incremental step sizes and mean single motor unit amplitudes. The compound response and mean step size were then combined to estimate MUNE (see Section 2.5). Example recruitment curves for OPM-MMG and EMG are shown at the bottom.

After filtering, stimulus-locked segmentation was performed using the hardware triggers generated by the stimulation system (see Section 2.2). Each electrical stimulation delivered to the ulnar nerve produced a synchronized trigger recorded in both acquisition systems, which served as the temporal reference for aligning the EMG and OPM-MMG signals. Continuous recordings were segmented into stimulus-centered epochs relative to these trigger events. For each trigger, an evoked-response window (or a signal window) was defined starting 5.5 ms after stimulus onset with a duration of 15 ms, corresponding to the expected latency and duration of the electrically evoked ADM response. To estimate baseline noise, a 15 ms noise window was extracted 150 ms prior to the stimulus trigger, ensuring that the window was free of stimulus-related activity. The stimulation intensity associated with each trigger was stored during acquisition and used to reconstruct the recruitment curves across stimulation levels. These signal and noise windows were subsequently used for signal quality assessment and amplitude quantification during MUNE calculation (Figure 2, Step 3).

After visual inspection, all OPM-MMG-signals of sensors positioned more than 2 cm from the skin surface (n = 12, see Figure1A) were excluded from further analysis, as they did not exhibit a clear stimulus to evoked response relationship and failed to yield a meaningful compound muscle action field to single motor unit field (CMAF/SMUF) ratio (see Figure 2, Step 2). Next, the filtered OPM-MMG and EMG data were segmented and used for the subsequent MUNE computation step, as described in detail in the following section.

### 2.5. MUNE calculation

As outlined above, several MUNE approaches have been proposed that differ primarily in how single motor unit contributions are estimated from compound muscle responses. The present analysis follows the principle of classical incremental stimulation MUNE, in which the average single motor unit amplitude is estimated from recruitment increments of the compound response and MUNE is calculated as the ratio between the supramaximal compound response and the mean single motor unit contribution (McComas et al., 1971). In contrast to the original incremental implementations, which typically rely on manual identification of a limited number of recruitment steps, the present approach derives recruitment increments from densely sampled stimulus–response curves that construct a whole MU pool and applies automated step-size filtering to obtain a representative estimate of single motor unit amplitude. This procedure preserves the incremental MUNE framework while differing from classical implementations in several aspects: recruitment increments are derived from densely sampled stimulus–response curves, the identification of increments is performed automatically rather than by manual selection, and the representative single motor unit amplitude is estimated from the filtered distribution of recruitment step sizes rather than from a small manually selected set of increments.

Recruitment analysis assumes that repeated responses at a given stimulation intensity are stable and reliable; signal stationarity and measurement consistency must first be ensured before constructing recruitment curves. Accordingly, the analysis pipeline first evaluates signal reliability and subsequently derives recruitment, incremental amplitudes, and MUNE from quality-controlled data (Figure 3). The same processing steps were applied to both OPM-MMG and EMG electrodes.

**Figure 3:**
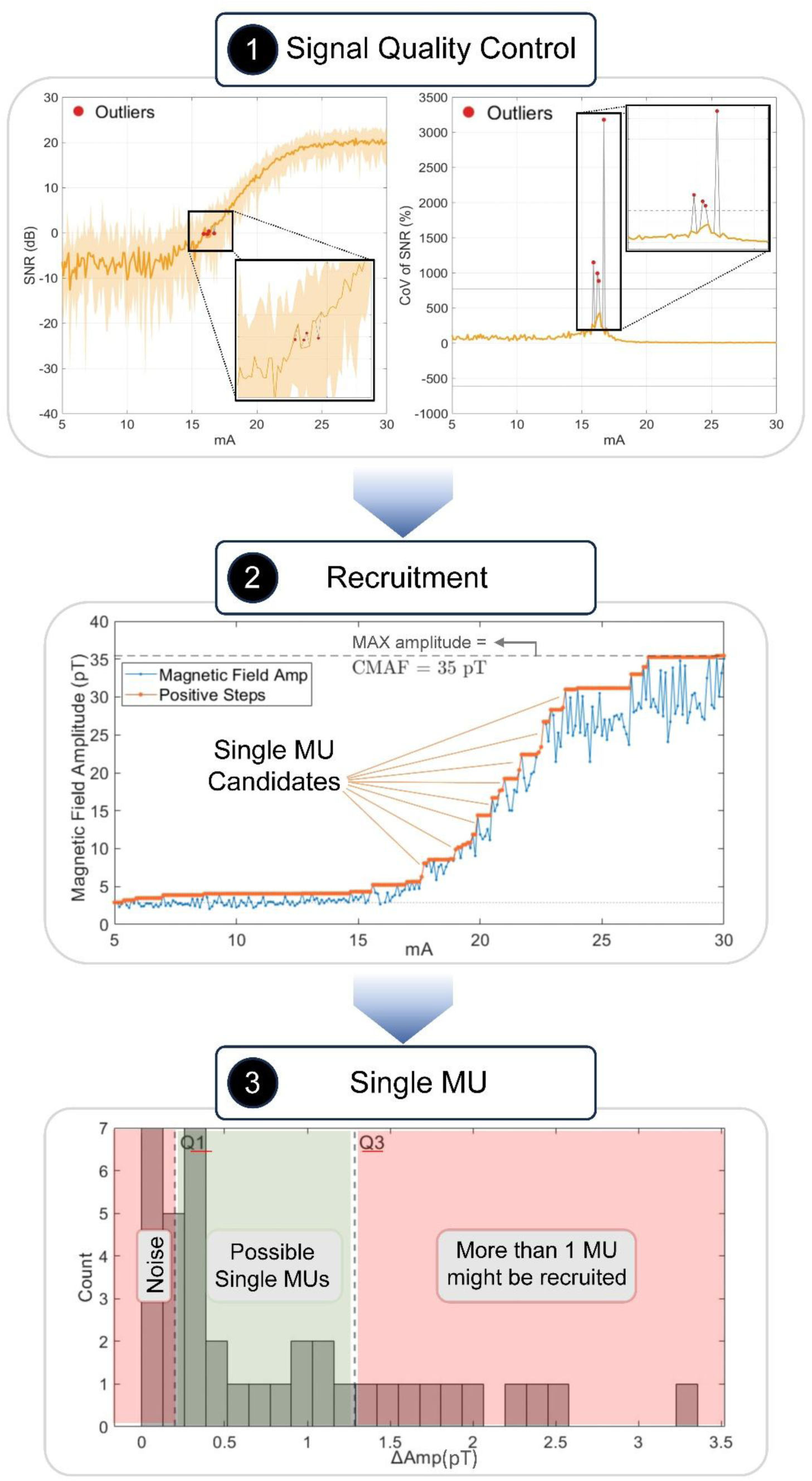
Stepwise MUNE analysis pipeline of an OPM-MMG sensor starting from signal-quality control to the final selection of single motor unit contributions: **Step 1** shows automated signal-quality screening based on SNR variability across stimulation intensity, where stimulation steps exhibiting excessive SNR-CV are identified as outliers and interpolated to ensure stable recruitment estimation. Red dots highlight outliers at specific stimulation levels. **Step 2** depicts the construction of the recruitment curve from peak-to-peak evoked response amplitudes, its transformation into a non-decreasing curve with positive steps, and the identification of candidate single motor unit fields. The supramaximal compound muscle action field is defined as the plateau amplitude of the recruitment curve. **Step 3** shows the distribution of recruitment step sizes, where steps within the interquartile range (Q1–Q3) are retained as physiologically plausible single motor unit contributions, while smaller steps are attributed to noise, and larger steps are likely due to the recruitment of multiple motor units.

First, an automated step-wise signal-quality screening was performed (Figure 3, Step 1). For each stimulation intensity and repetition, the single-stimulus SNR was computed using an evoked-response window and a matched noise window (Figure 2, Step 3). SNR variability across repetitions was summarized at each stimulation intensity using the coefficient of variation (CV), yielding one SNR-CV value per stimulation level. Because repeated stimuli at a fixed intensity are expected to reflect the same recruited MU population, high SNR variability (exceeding ±3 standard deviations of the SNR-CV distribution across stimulation intensities) primarily reflects transient measurement instability (e.g., motion or environmental interference) rather than physiological effects. Stimulation responses with SNR-CV values exceeding the mean ±3 standard deviations of the SNR-CV distribution across stimulation intensities were classified as unstable and interpolated.

Second, the evoked response amplitude was quantified from the stimulus-locked waveform segments using the peak-to-peak amplitude,

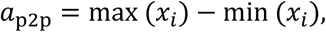

where 𝑥_𝑖_ denotes the evoked muscle response window.

Third, given that the evoked response amplitudes physiologically vary over time (i.e., the same stimulation does not necessarily yield the same signal amplitude), we transformed the recruitment curve into a non-decreasing curve with only positive steps, which then allows MUNE calculation (Figure 3, Step 2). The supramaximal compound response (CMAP/CMAF) amplitude was defined as the difference between the maximum value of the recruitment curve and its baseline.

Fourth, we estimated the average amplitude of a single MU. The distribution of recruitment step sizes was summarized (Figure 3, Step 3), where candidate single motor units were defined as positive consecutive differences in the recruitment curve. Step sizes falling below the first quartile (Q1) were considered likely to reflect noise-related fluctuations or subthreshold variability, whereas step sizes above the third quartile (Q3) were assumed to reflect the near-simultaneous recruitment of multiple motor units rather than a single unit. Accordingly, a representative single motor unit amplitude was estimated by retaining only step sizes within the interquartile range (Q1–Q3). MUNE was then computed for each of the 20 repetitions as

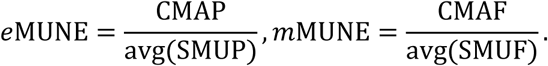

Finally, as an additional step, all MUNE values were screened for outliers using the interquartile range criterion (±1.5×IQR). Flagged values were retained as missing for downstream analyses rather than being imputed, resulting in a minimum sample size of 18 and a maximum of 20 accepted MUNE values.

### 2.5. Statistical analysis

All analyses were performed on each subject, modality, and sensor/electrode. MUNE was computed independently for each stimulation repetition (min n=18, max n=20 per sensor or electrode), yielding a distribution of MUNE per subject and sensor/electrode. These distributions were summarized using the median and interquartile range (1st–3rd quartile). Within-measure variability was quantified using the coefficient of variation, calculated across repetitions for each MUNE, CMAF/CMAP, and SMUP/SMUF. Signal quality was summarized using median SNR and corresponding variability measures.

To evaluate whether cross-modality differences in MUNE were primarily attributable to differences in signal quality, statistical comparisons were performed before and after SNR matching. Repetition-wise MUNE values obtained from OPM-MMG sensors and EMG electrodes were treated as paired observations. Each OPM-MMG sensor was paired with the closest EMG electrode. The normality of paired differences was assessed using the Lilliefors test. As the paired differences deviated from normality, non-parametric Wilcoxon signed-rank tests were used to assess differences between conditions. Two comparisons were evaluated: OPM-MMG versus EMG-derived MUNE before SNR matching (described below), OPM-MMG versus EMG-derived MUNE after SNR matching. To account for multiple comparisons, p-values were adjusted using the Benjamini–Hochberg false discovery rate procedure.

### 2.6. Influence of noise on MUNE

To investigate the effect of SNR on MUNE, we matched SNR between EMG and OPM-MMG recordings. For each subject and corresponding OPM–EMG sensor pair, SNR was estimated from stimulation-locked response segments using the same segmentation procedure employed for the main analysis. Signal amplitude was quantified using noise-corrected root-mean-square (RMS) values, where the baseline noise level was estimated from pre-stimulus windows.

The mean OPM-MMG SNR at each stimulation step served as the target SNR for the corresponding EMG electrode. To achieve this target SNR, additive noise was injected into the EMG signal to match the OPM-MMG SNR. Specifically, the required noise amplitude was determined from the relationship between the noise-corrected signal RMS and the desired noise RMS corresponding to the target SNR. Band-limited (10-185 Hz) white Gaussian noise (WGN) was generated and added to the EMG signal segments. This bandwidth was selected to match the effective spectral range of the filtered EMG/MMG signals. The additive noise increased the baseline noise level of the EMG recordings while preserving the underlying evoked response morphology. As a result, the SNR of the augmented EMG signals was reduced to levels comparable to those observed in OPM-MMG recordings, enabling a controlled comparison of MUNE estimates across modalities under matched signal-quality conditions.

## 3. Results

### 3.1. Sigmoidal Recruitment Curve

As stimulation intensity increased, OPM-MMG and EMG signals exhibited the characteristic sigmoidal recruitment behavior, with discrete step-like increments at lower intensities and reaching a plateau at higher currents, consistent with motor unit recruitment dynamics (Figure 4). Recruitment curves derived from OPM-MMG closely matched those from EMG, demonstrating that OPM-MMG measured the same underlying recruitment process across the full stimulation range.

**Figure 4:**
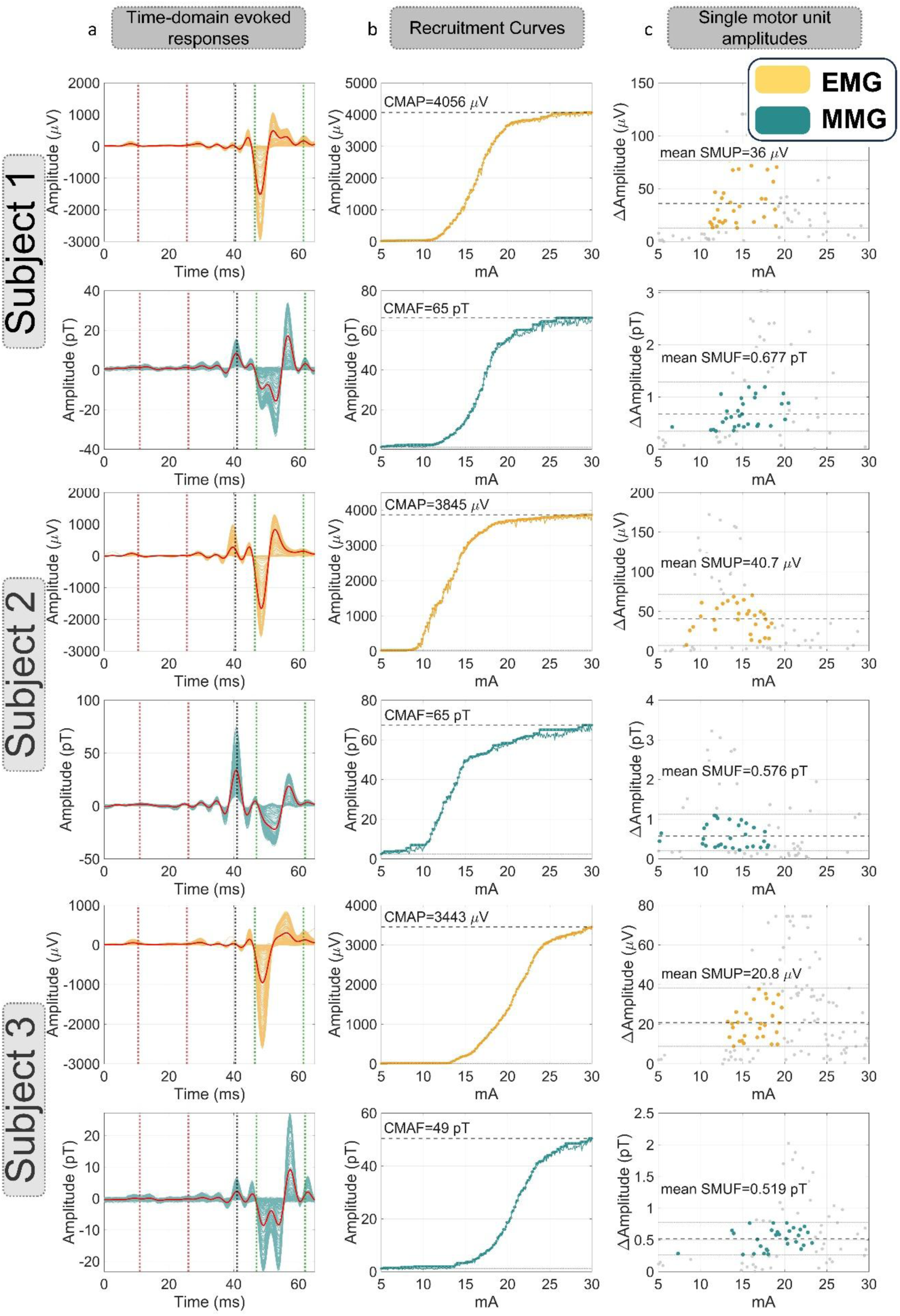
Snippets of ADM signal under all stimulation intensities across EMG and closest OPM-MMG sensors (yellow: EMG; teal: OPM-MMG). **(a)** Time-domain evoked responses showing stimulus-locked EMG potentials (µV) and OPM-MMG magnetic field signals (pT) across repeated stimulations, illustrating waveform morphology and temporal consistency (red line indicates the average signal across stimulus waveforms). **(b)** Recruitment curves depicting compound response amplitudes as a function of stimulation current, yielding CMAP for EMG and CMAF for OPM-MMG, with a characteristic sigmoid curve. **(c)** Single motor unit amplitudes plotted against stimulation intensity, showing discrete SMUP (EMG) and SMUF (OPM-MMG) estimates and their variability across recruitment.

### 3.2. Signal quality

Signal quality and motor unit metrics for all subjects are summarized in Table 1, reported separately for OPM-based MMG and surface EMG electrodes. Across subjects, OPM sensors exhibited lower median SNR values than EMG electrodes, with median OPM SNRs ranging from approximately 18 to 27 dB, depending on subject and sensor, whereas median EMG SNRs were typically around 29–31 dB. The variability of median SNRs across repetitions was modest for both modalities within-session, with coefficients of variation generally between ∼8–17% for OPM sensors and ∼3–10% for EMG electrodes, indicating stable signal quality within each modality despite lower absolute SNR for OPM. One EMG electrode (EMG-04, Table 1) from Subject 1 elicited low SNR values across repetitions and was thus excluded from further analysis.

**Table 1.**
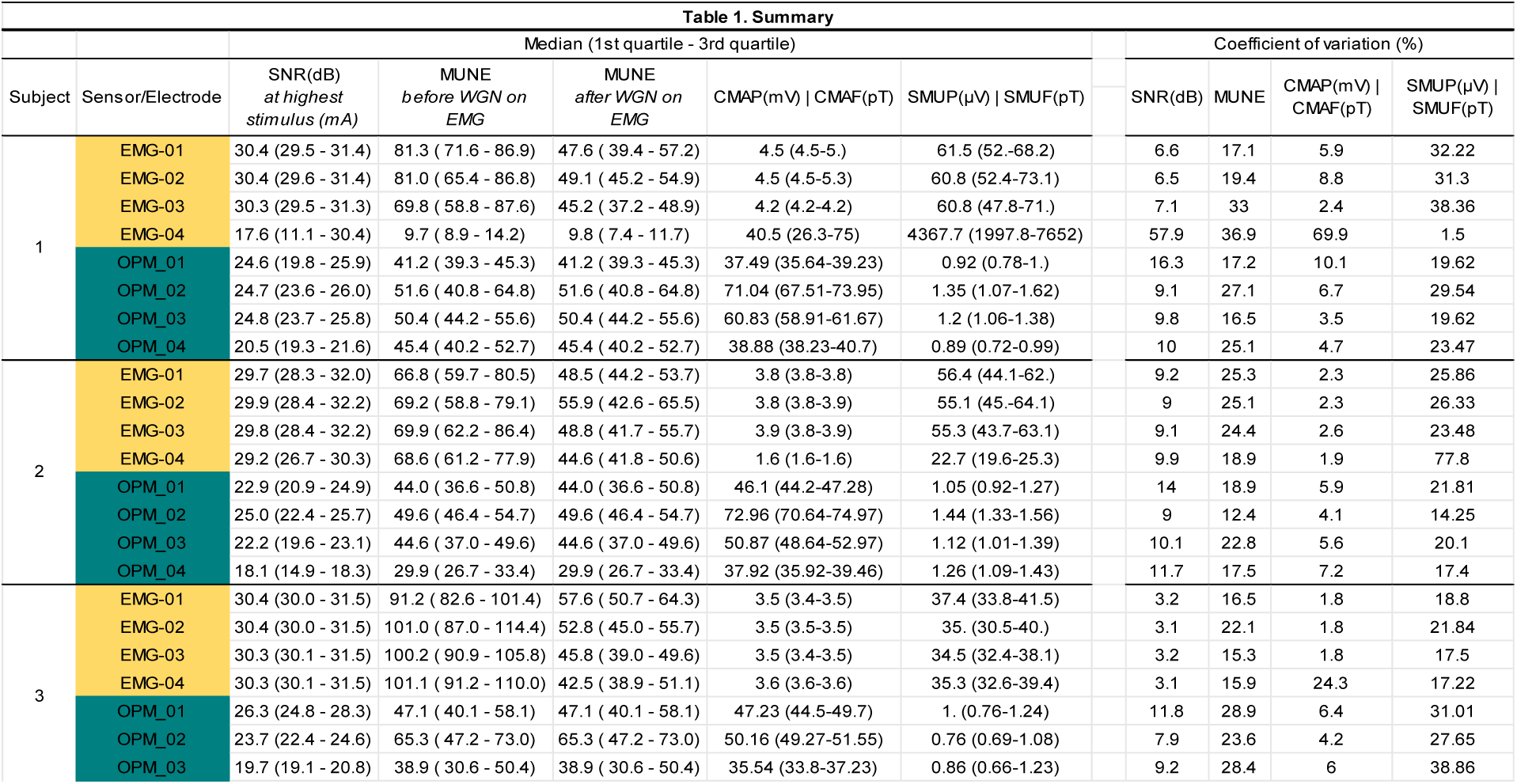
Signal quality and motor unit metrics across subjects and sensors. Median values (1st–3rd quartile) are shown for signal-to-noise ratio (SNR), motor unit number estimation (MUNE), MUNE results with added white Gaussian noise (WGN) applied to EMG only, compound muscle action potential/field amplitude (CMAP for EMG in mV; CMAF for OPM in pT), and single motor unit potential/field amplitude (SMUP for EMG in µV; SMUF for OPM in pT). Coefficients of variation (CV, %) are provided separately. Importantly, the column “MUNE with WGN on EMG” reflects MUNE values after synthetic white Gaussian noise was added exclusively to the EMG signals to match lower SNR conditions; no noise was added to OPM-MMG signals for this analysis. The fourth electrode of Subject 1 (EMG-04, most distal position) exhibited low SNR values and was excluded from further analysis.

### 3.3. Motor unit number estimation

mMUNE and eMUNE estimates differed and showed moderate variability across sensors. mMUNE values were significantly lower than eMUNE values. Across all paired repetition-wise estimates, EMG electrodes yielded a median MUNE of 78.81, whereas OPM-MMG sensors yielded a median MUNE of 45.41. This difference between modalities was statistically significant (Wilcoxon signed-rank test, FDR-corrected p <0.001). mMUNE ranged from ∼30 to ∼65, with coefficients of variation generally between ∼12–31% across sensors. In contrast, eMUNE values ranged from ∼69 to ∼101 across repetitions and coefficients of variation typically between ∼15% and ∼37% across sensors. Within each subject, the relative spread of mMUNE and eMUNE values across sensors was similar, indicating that differences between modalities were not driven by increased variability (Figure 5).

**Figure 5:**
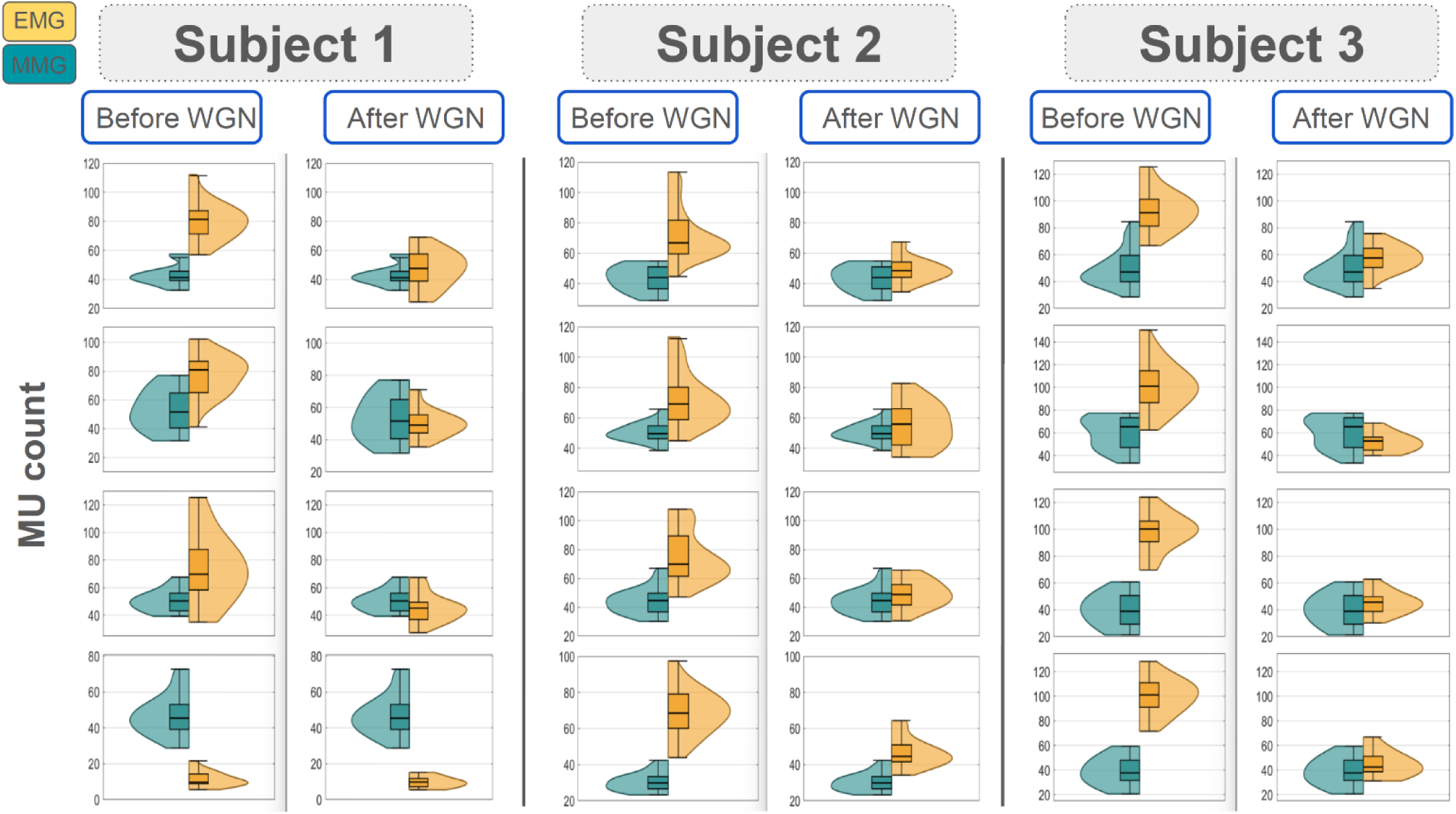
Comparison of 4 OPM-MMG sensors and the 4 EMG electrodes that are positioned right next to each other during the experiment. For each subject n= (min=18, max=20) MUNE samples obtained without and with added white Gaussian noise (WGN) to the EMG signal. WGN was added to EMG signals to reduce the SNR to levels comparable to those of OPM-MMG. After the WGN added and SNRs were matched, the difference between modalities was no longer statistically significant (Wilcoxon signed-rank test, FDR-corrected p = 0.55). These results indicate that the observed cross-modality differences in MUNE primarily reflect differences in signal-to-noise ratio rather than fundamental differences in the measurement modality.

### 3.4. Evoked compound motor unit responses

At the compound motor unit response level, CMAF amplitudes ranged from ∼33.9 to 73 pT and showed slightly higher, yet still moderate, variability, with coefficients of variation typically between 4–11%. Corresponding CMAP amplitudes ranged approximately from ∼3.4 to 6.4 mV across subjects and sensors and showed low variability, with coefficients of variation mostly below ∼10%, reflecting stable maximal responses.

### 3.5. Evoked single motor unit responses

Single motor unit responses exhibited greater variability than compound responses. Median SMUF amplitudes ranged from ∼0.7 to 1.4 pT, and showed variability with coefficients of variation typically between ∼14–39 %. SMUP amplitudes ranged approximately from ∼20 to 85 µV across subjects and sensors, with coefficients of variation between ∼18–78%.

### 3.6. Influence of noise on MUNE

To test whether EMG-derived MUNE values were significantly higher than MMG-derived MUNE values due to the higher SNR of EMG, we matched the SNR across modalities by augmenting the EMG data with additive noise. After applying the SNR matching procedure, the difference between modalities vanished (Figure 5). Under matched SNR conditions, EMG-derived MUNE estimates (median = 47.04) closely matched OPM-MMG-derived estimates (median = 45.41), and no statistically significant difference between modalities was observed (Wilcoxon signed-rank test, FDR-corrected p = 0.55).

## 4. Discussion

Here, we demonstrate that contactless mMUNE can be performed using OPM-MMG within a distance of ∼2 cm. We found that OPM-MMG shows the same characteristic sigmoidal recruitment behavior as EMG (Figures 2 and 3). Median mMUNE was consistently lower across OPM-MMG sensors than median eMUNE across EMG electrodes by approximately 40%. This difference was accompanied by an approximately 25% lower SNR of OPM-MMG recordings (Table 1 and Figure 5). Equalizing SNR between modalities, we identified this lower SNR in OPM-MMG as the primary reason for the reduced mMUNE compared to eMUNE.

The mMUNE approach enabled us to quantify detailed reference values of MMG-SMUF and MMG-CMAF measurements. SMUF measurements ranged from 0.7 to 1.4 pT, whereas the CMAF values ranged from 34 to 73 pT (Figure 3, Table 1). These SMUF values can be interpreted in the context of prior SQUID-MMG and OPM-based MMG studies that attempted to estimate SMUF during voluntary activity. As such, (Masuda et al., 1999) reported SMUF amplitudes of 1–2 pT using SQUID-MMG in the vastus medialis and vastus lateralis during voluntary knee extension. In contrast, we observed lower median SMUF amplitudes of approximately 0.7 pT in the ADM muscle. This difference is likely attributable to anatomical and physiological factors. The quadriceps muscles are substantially larger than the ADM muscle and typically contain motor units comprising a greater number of muscle fibers, which is likely to generate larger electromagnetic signals (Purves, 2001). Another factor that may contribute to differences in measured amplitudes is the sensor–muscle distance, as Masuda et al. reported SQUID sensors positioned approximately 40 mm from the current source, whereas in the present study OPM sensors were positioned at a range of approximately 5-20 mm from the skin. Because magnetic field strength decreases rapidly with increasing source–sensor distance, such methodological differences may also influence the observed amplitudes. Similarly, a recent study on single motor unit decomposition of MMG recordings reported single ADM unit amplitudes ranging from 0.75 to 12 pT across sensors (Noury et al., 2025). However, their paradigm did not ensure a constant sensor-to-skin distance, and in some cases, they reported distances of about 1 to 2 mm. Considering this variable and shorter distance, their findings are generally in line with the values observed in our study. In addition, Masuda et al. employed a feedback-controlled voluntary force output of approximately 10–20% of the subject’s maximum voluntary activation (MVC), which is expected to involve more than a single motor unit. In contrast, the evoked paradigm employed here enables a more precise quantification of the SMUF.

A related and recent comparison to our study’s investigation on the single motor unit field is by (Otani et al., 2025), who also employed electrically evoked activation but reported source-space current moments (0.44–9.05 nA·m in healthy participants) in the abductor pollicis brevis rather than sensor-space magnetic field amplitudes. In their study, 12 single motor units were identified across 8 healthy participants using an all-or-none stimulation paradigm, with each unit reconstructed from multiple repetitions (5–500 responses) to improve SNR. In contrast, Masuda et al. estimated larger dipole moments (23.9–114.3 nA·m) in the vastus medialis and lateralis using equivalent dipole fitting of the final propagation peak. The smaller current moments reported by Otani et al. are consistent with both the smaller muscle studied and the different current component quantified, as Otani analyzed an early neuromuscular junction–related current rather than the later tendon-associated dipole peak analyzed by Masuda. Although direct numerical comparison with our study is limited because of the different physical units (pT vs. nA·m), both prior studies focused on isolated threshold-level motor units. In contrast, the incremental mMUNE approach applied here systematically recruits the whole motor unit pool and extracts a distribution of SMUF amplitudes across recruitment levels. This provides a population-level characterization of motor unit magnetic field magnitudes in the ADM rather than a single-unit estimate under voluntary or minimal activation conditions.

As stated above, SNR was a key determinant of MUNE in the present study, with lower OPM-MMG SNR resulting in lower mMUNE values. We attribute this low SNR not only to OPM sensor performance but also to the distance from the signal source. Previous work from our group examined the effect of sensor-to-source distance on MMG and reported that SNR decreased from ∼2.10 at 1 cm to ∼1.06 at 2 cm, and to <0.5 beyond 3 cm (Yang et al., 2025a), i.e., the signal became indistinguishable from noise beyond ∼2 cm. Direct comparison is constrained by methodological differences in the muscle studied, the type of activity, and the computation of SNR. As such, (Yang et al., 2025a) reported linear SNR ratios and likely obtained lower signals from voluntary contractions of a smaller muscle, whereas the present study achieved higher logarithmic SNR through electrical stimulation and strict stimulus-locking in the smaller ADM muscle.

Beyond these distance effects on SNR, OPM-MMG SNR could also be influenced by the size of the muscle studied. In another recent work of our group, (Baier et al., 2025) recorded OPM-MMG from the comparably larger right biceps brachii muscle during steady and force-controlled isometric contractions at 20%, 40%, and 60% MVC in 20 healthy adults, reporting force-dependent SNR increases from 11.10 ± 4.13 dB (20% MVC) to 16.67 ± 4.70 dB (40% MVC) and 21.37 ± 5.24 dB (60% MVC). In contrast, the present study quantified SNR under electrically evoked, stimulus-locked conditions, defined as the RMS ratio of the supramaximal evoked response at the recruitment plateau to its baseline noise window, yielding values of 18–27 dB. Thus, SNR values reported here are comparable to those reported at 60% MVC by Baier et al., which likely reflects the smaller size of the ADM muscle as compared to the biceps brachii muscle. Additionally, in the present study, OPM-MMG sensors were positioned slightly farther from the geometric center of the magnetically shielded room than in Baier et al., because the ADM muscle is more distal than the biceps brachii muscle. (Voigt et al., 2013) showed in an AK3b-type magnetically shielded room that residual fields (∼50 nT after delivery) and gradients (<30 pT/mm) vary spatially and are lowest within a defined central volume after optimized degaussing. Because vibration-induced magnetic noise scales with the local field gradient, small differences in sensor position within the room can directly affect the baseline noise floor.

### 4.1 Strengths and limitations

The primary strength of this study lies in the large number of repeated stimulations per subject, enabling a large sample characterization of within-session stability of mMUNE and eMUNE across sensors. This increased statistical power for detecting single motor unit amplitudes and obtaining a more reliable distribution of the ADM MU pool within a single session. Another key strength is the simultaneous OPM-MMG and EMG recording, which ensured that both modalities recorded the same underlying recruitment processes, enabling a well-controlled, direct comparison.

Several limitations should be acknowledged. First, OPM-MMG sensors require a magnetically shielded environment, which naturally limits the portability of potential mMUNE applications. Since MUNE has been shown to be an important biomarker for certain neuromuscular diseases in previous studies (Gooch et al., 2014; Shefner et al., 2011; Swoboda et al., 2005) and for clinical translation of the mMUNE, a portable magnetic shield would be essential. Considering this, there were recent significant developments to make OPM-MMG more portable. (Nordenström et al., 2024) demonstrated the feasibility of OPM-MMG in a commercially available mobile magnetic shield (Twinleaf MS-2), where they recorded electrically evoked ADM responses to ulnar nerve stimulation using a triaxial zero-field OPM and directly compared the resulting waveforms to recordings obtained in the Berlin magnetically shielded room (BMSR-2). Although the mobile shield exhibited substantially higher baseline noise spectral densities (≈70–300 fT/√Hz) compared to the heavily shielded BMSR-2 (<20 fT/√Hz), averaged evoked MMG waveforms remained comparable, supporting the practical feasibility of portable OPM-MMG despite elevated environmental noise. Second, the present study focused on a small number of healthy participants and was designed to establish methodological feasibility. While this approach is appropriate for a proof of concept, future work should evaluate mMUNE in larger, more diverse cohorts, including patient populations where mMUNE is clinically informative. Third, the strong dependence on a close sensor-to-skin distance currently limits the signal quality of OPM-MMG sensors, underscoring the importance of optimized sensor placement and noise-floor reduction strategies. The final limitation is the lack of reliability assessment between measurements on the same subject at different times. This study mainly focused on within-session reliability and given that OPM-MMG heavily relies on sensor placement relative to the muscle, it will be important to investigate how reproducible results are across sessions. This is particularly relevant for monitoring a decrease in motor unit count in patients with ongoing denervation.

### 4.2 Conclusion

To conclude, our findings demonstrate that OPM-MMG enables contactless motor unit number estimation (MUNE).

## Acknowledgment and funding

The authors would like to thank all participants for their time, dedication, and contribution to this study. The funding sources were not involved in the study design, data collection, analysis, interpretation of data, or in the writing of the manuscript.

B.S., T.K., and J.M. are supported by the Deutsche Forschungsgemeinschaft (DFG, German Research Foundation) through the priority program SPP 2311 (Grant ID: 548605919). T.K., J.M., and O.R. are supported by the European Research Council (ERC) through the ERC Advanced Grant “qMOTION” (Grant ID: 101055186). J.M. and N.N. are supported by the Bundesministerium für Bildung und Forschung (BMBF) through the Future Cluster “QSens” (Grant ID: 03ZU2110FD). B.S. and J.M. are supported by the Deutsches Zentrum für Luft- und Raumfahrt (DLR) (Grant ID: 50BM2534B).

## References

Baier, L., Brümmer, T., Senay, B., Siegel, M., Keleş, A.D., Röhrle, O., Klotz, T., Noury, N., Marquetand, J., 2025. Contactless measurement of muscle fiber conduction velocity—a novel approach using optically pumped magnetometers. J. Neural Eng. 22, 026058. 10.1088/1741-2552/adc83b

Boe, S.G., Stashuk, D.W., Doherty, T.J., 2004. Motor unit number estimation by decomposition-enhanced spike-triggered averaging: Control data, test–retest reliability, and contractile level effects. Muscle & Nerve 29, 693–699. 10.1002/mus.20031

Bostock, H., 2016. Estimating motor unit numbers from a CMAP scan. Muscle & Nerve 53, 889–896. 10.1002/mus.24945

Boto, E., Meyer, S.S., Shah, V., Alem, O., Knappe, S., Kruger, P., Fromhold, T.M., Lim, M., Glover, P.M., Morris, P.G., Bowtell, R., Barnes, G.R., Brookes, M.J., 2017. A new generation of magnetoencephalography: Room temperature measurements using optically-pumped magnetometers. NeuroImage 149, 404–414. 10.1016/j.neuroimage.2017.01.034

Broser, P.J., Marquetand, J., Middelmann, T., Sometti, D., Braun, C., 2021. Investigation of the temporal and spatial dynamics of muscular action potentials through optically pumped magnetometers. J Electromyogr Kinesiol 59, 102571. 10.1016/j.jelekin.2021.102571

Brown, W.F., Strong, M.J., Snow, R., 1988. Methods for estimating numbers of motor units in biceps-brachialis muscles and losses of motor units with aging. Muscle & Nerve 11, 423–432. 10.1002/mus.880110503

Chen, M., Lu, Z., Zong, Y., Li, X., Zhou, P., 2023. A Novel Analysis of Compound Muscle Action Potential Scan: Staircase Function Fitting and StairFit Motor Unit Number Estimation. IEEE Journal of Biomedical and Health Informatics 27, 1579–1587. 10.1109/JBHI.2022.3229211

Cohen, D., Givler, E., 1972. Magnetomyography: magnetic fields around the human body produced by skeletal muscles. Applied Physics Letters 21, 114–116. 10.1063/1.1654294

Doherty, T.J., Brown, W.F., 1993. The estimated numbers and relative sizes of thenar motor units as selected by multiple point stimulation in young and older adults. Muscle & Nerve 16, 355–366. 10.1002/mus.880160404

Doherty, T.J., Stashuk, D.W., 2003. Decomposition-based quantitative electromyography: Methods and initial normative data in five muscles. Muscle & Nerve 28, 204–211. 10.1002/mus.10427

Farina, D., Holobar, A., Merletti, R., Enoka, R.M., 2010. Decoding the neural drive to muscles from the surface electromyogram. Clinical Neurophysiology 121, 1616–1623. 10.1016/j.clinph.2009.10.040

Fuglsang-Frederiksen, A., 2006. The role of different EMG methods in evaluating myopathy. Clinical Neurophysiology 117, 1173–1189. 10.1016/j.clinph.2005.12.018

Gooch, C.L., Doherty, T.J., Chan, K.M., Bromberg, M.B., Lewis, R.A., Stashuk, D.W., Berger, M.J., Andary, M.T., Daube, J.R., 2014. Motor unit number estimation: A technology and literature review. Muscle & Nerve 50, 884–893. 10.1002/mus.24442

Gooch, C.L., Pullman, S.L., Shungu, D.C., Uluğ, A.M., Chane, S., Gordon, P.H., Tang, M.X., Mao, X., Rowland, L.P., Mitsumoto, H., 2009. Motor unit number estimation (MUNE) in diseases of the motor neuron: utility and comparative analysis in a multimodal biomarker study. Suppl Clin Neurophysiol 60, 153–162. 10.1016/s1567-424x(08)00015-9

Hansen, S., Ballantyne, J.P., 1978. A quantitative electrophysiological study of motor neurone disease. J Neurol Neurosurg Psychiatry 41, 773–783. 10.1136/jnnp.41.9.773

Kaya, R.D., Hoffman, R.L., Clark, B.C., 2014. Reliability of a modified motor unit number index (MUNIX) technique. Journal of Electromyography and Kinesiology 24, 18–24. 10.1016/j.jelekin.2013.10.005

Kominis, I.K., Kornack, T.W., Allred, J.C., Romalis, M.V., 2003. A subfemtotesla multichannel atomic magnetometer. Nature 422, 596–599. 10.1038/nature01484

Marquetand, J., Middelmann, T., Dax, J., Baek, S., Sometti, D., Grimm, A., Lerche, H., Martin, P., Kronlage, C., Siegel, M., Braun, C., Broser, P., 2021. Optically pumped magnetometers reveal fasciculations non-invasively. Clin Neurophysiol 132, 2681–2684. 10.1016/j.clinph.2021.06.009

Masuda, T., Endo, H., Takeda, T., 1999. Magnetic fields produced by single motor units in human skeletal muscles. Clin Neurophysiol 110, 384–389. 10.1016/s1388-2457(98)00021-2

McComas, A.J., Fawcett, P.R.W., Campbell, M.J., Sica, R.E.P., 1971. Electrophysiological estimation of the number of motor units within a human muscle. Journal of Neurology, Neurosurgery & Psychiatry 34, 121–131. 10.1136/jnnp.34.2.121

McNeil, C.J., Doherty, T.J., Stashuk, D.W., Rice, C.L., 2005. Motor unit number estimates in the tibialis anterior muscle of young, old, and very old men. Muscle & Nerve 31, 461–467. 10.1002/mus.20276

Nordenström, S., Lebedev, V., Hartwig, S., Kruse, M., Marquetand, J., Broser, P., Middelmann, T., 2024. Feasibility of magnetomyography with optically pumped magnetometers in a mobile magnetic shield. Sci Rep 14, 18960. 10.1038/s41598-024-69829-y

Noury, N., Marquetand, J., Hartwig, S., Middelmann, T., Broser, P., Siegel, M., 2025. Detecting single motor-unit activity in magnetomyography. J. Neural Eng. 22, 046023. 10.1088/1741-2552/adeaeb

Oostenveld, R., Fries, P., Maris, E., Schoffelen, J.-M., 2011. FieldTrip: Open source software for advanced analysis of MEG, EEG, and invasive electrophysiological data. Comput Intell Neurosci 2011, 156869. 10.1155/2011/156869

Otani, T., Akaza, M., Kawabata, S., Natsui, H., Watanabe, T., Miyano, Y., Hanazawa, R., Adachi, Y., Sekihara, K., Kanouchi, T., Yokota, T., 2025. Magnetomyographic evaluation of motor unit size that is robust to changes in distance to sensor. Brain Commun 7, fcaf294. 10.1093/braincomms/fcaf294

Power, G.A., Dalton, B.H., Rice, C.L., 2013. Human neuromuscular structure and function in old age: A brief review. J Sport Health Sci 2, 215–226. 10.1016/j.jshs.2013.07.001

Purves, 2001. Neuroscience. Sinauer Associates Inc., Sunderland, Mass.

Shefner, J.M., Watson, M.L., Simionescu, L., Caress, J.B., Burns, T.M., Maragakis, N.J., Benatar, M., David, W.S., Sharma, K.R., Rutkove, S.B., 2011. Multipoint incremental motor unit number estimation as an outcome measure in ALS. Neurology 77, 235–241. 10.1212/WNL.0b013e318225aabf

Sherrington, C.S., 1919. Mammalian physiology; a course of practical exercises. Clarendon Press, Oxford. 10.5962/bhl.title.20939

Swoboda, K.J., Prior, T.W., Scott, C.B., McNaught, T.P., Wride, M.C., Reyna, S.P., Bromberg, M.B., 2005. Natural history of denervation in SMA: Relation to age, SMN2 copy number, and function. Annals of Neurology 57, 704–712. 10.1002/ana.20473

Voigt, J., Knappe-Grüneberg, S., Schnabel, A., Körber, R., Burghoff, M., 2013. Measures to reduce the residual field and field gradient inside a magnetically shielded room by a factor of more than 10. Metrology and Measurement Systems Vol. 20.

Yang, H., Klotz, T., Gizzi, L., Lu, H., Monittola, G., Schneider, U., Siegel, M., Marquetand, J., 2025a. The effect of sensor-to-source distance on magnetic neuromuscular signals. Sci Rep 15, 20225. 10.1038/s41598-025-06545-1

Yang, H., Senay, B., Sorrentino, C., Bouraima, F., Siegel, M., Marquetand, J., 2025b. Real-time distance monitoring in magnetomyography. J. Neural Eng. 22, 066002. 10.1088/1741-2552/ae1874

